# A Computational Workflow for Analysis of Missense Mutations in Precision Oncology

**DOI:** 10.1101/2024.06.08.598055

**Authors:** Rayyan Tariq Khan, Petra Pokorna, Jan Stourac, Simeon Borko, Ihor Arefiev, Joan Planas-Iglesias, Adam Dobias, Gaspar Pinto, Veronika Szotkowska, Jaroslav Sterba, Ondrej Slaby, Jiri Damborsky, Stanislav Mazurenko, David Bednar

## Abstract

Every year, more than 19 million cancer cases are diagnosed, and this number continues to increase annually. Since standard treatment options have varying success rates for different types of cancer, understanding the biology of an individual’s tumour becomes crucial, especially for cases that are difficult to treat. Personalised high-throughput profiling, using next-generation sequencing, allows for a comprehensive examination of biopsy specimens. Furthermore, the widespread use of this technology has generated a wealth of information on cancer-specific gene alterations. However, there exists a significant gap between identified alterations and their proven impact on protein function. Here, we present a bioinformatics pipeline that enables fast analysis of a missense mutation’s effect on stability and function in known oncogenic proteins. This pipeline is coupled with a predictor that summarises the outputs of different tools used throughout the pipeline, providing a single probability score, achieving the balanced accuracy above 86%. The pipeline incorporates a virtual screening method to suggest potential FDA/EMA-approved drugs to be considered for treatment. We showcase three case studies to demonstrate the timely utility of this pipeline. To facilitate access and analysis of cancer-related mutations, we have packaged the pipeline as a web server, which is freely available at https://loschmidt.chemi.muni.cz/predictonco/.

## Introduction

More than 19 million cancer cases were diagnosed in 2020 (Cancer Today, 2020) with a projected load of 28.4 million cases in 2040 (Sung et al., 2021). The three traditionally used approaches to treat cancer, namely chemotherapy, surgery, and radiotherapy, generally result in higher mortality rates compared to the less adopted precision medicine-based techniques (Krzyszczyk et al. 2018). Next-generation sequencing technologies form the basis of precision oncology and can help generate a large amount of transcriptomic and genomic data. On the other hand, these technologies often do not provide clinically actionable data. This leads to a divide between generation of the said data and their utility, as mutants with unknown effects are often found during clinical testing (Buzdin et al., 2021).

There are not many tools that can help bridge the gap between data generation and creation of actionable insights. Swiss-PO, an online tool, allows for mapping experimentally determined mutations on a curated list of 50 genes and their various associated 3D structures. It also allows users to visualise multiple molecular interactions; however, it leaves it to the user to intuitively assess the structural implications of mutations that have not been experimentally determined (Krebs et al., 2021) and it can also not predict patient survival outcomes. PSnpBind, a database, catalogues changes to binding affinities of ligands due to binding site single-nucleotide polymorphisms (SNPs), however this database is limited to 26 human proteins and is limited to interactions between ligands and binding site residues (Ammar et al., 2022). We sought to overcome some of these limitations by creating a robust pipeline that can predict the effects of missense mutations, even for ones which are not experimentally determined, on cancer-related proteins.

The pipeline relies on advances in fast protein modelling, such as AlphaFold (Jumper et al., 2021), prediction of the effect of missense mutations on a protein structure (Bendl et al., 2014), and protein stability prediction (Blanco et al., 2018; Kellogg et al., 2010). This allows for a lot more information to be harvested from mutations identified by exome sequencing and can then be used for actionable decision making. Additionally, coupling fast ligand docking in proteins (Trott & Olson, 2009) and with the availability of multiple drug libraries online, such as ZINC (Irwin et al., 2012), it is possible to screen novel potential inhibitors for the mutated proteins.

As the interpretation of large-scale genomic and transcriptomic data is limited due to the need to utilise multiple computational tools, performing the aforementioned analysis on exome sequences can take time if done manually. After a cancer diagnosis, treatment is generally a race against time, and with the variable success rates of conventional “one size fits all” therapies, fast and accurate interpretation of molecular findings and assessment of their actionability are of vital importance, especially in difficult-to-treat cases. This is where an automated precision oncology approach will be most useful as it can optimise treatment strategies, improve outcomes and increase the quality of life for many patients (Lassen et al., 2021).

Here we introduce a bioinformatics pipeline for the analysis of the effect of mutations on stability and function in cancer-related proteins. The pipeline applies *in silico* methods of molecular modelling, structural bioinformatics and machine learning, and outputs actionable data which can be used for decision making. The coupled predictor produces a decision on the oncogenicity of the protein mutation by utilising the outputs derived at various stages of the pipeline. Moreover, we show the application of the pipeline on three use cases and highlight the importance of advanced bioinformatics in precision oncology.

## Methodology

### Manual Curation, Structure Repairs and Geometry Optimization

A list of 44 cancer-related proteins (including 1 isoform of a selected protein) were chosen as targets for the manual curation. The selection was based on the importance of the respective proteins for cancer diagnostics and, notably, in cancer treatment. The vast majority of curated proteins are either direct targets of therapeutic agents or, despite not being targets themselves, represent established predictive biomarkers for administering targeted treatments aimed at downstream members of the same pathway. Additionally, we included proteins that are frequently altered across various cancer types and are relevant to both diagnostics and cancer research (e.g., p53). The proteins with their various annotations are listed in the Supplementary material **[SI 1](https://docs.google.com/spreadsheets/d/1wuE5v_9v-_mS6be1go6eWHxTpCl2vfk3nP-mMJ6-f0Q/edit#gid=757186111)**.

The 44 protein sequences and their annotations were fetched from the UniProt database (The UniProt Consortium, 2023). In the case of KRAS, two isoforms are provided, including the canonical isoform and an isoform that is commonly utilized accross clinical databases of genetic variants. The essential residues were re-confirmed in the literature as well as in the Mechanism and Catalytic Site Atlas (M-CSA, Ribeiro et al., 2017) and the SWISS-PROT (Boeckmann, 2003) databases. For the purposes of this study, in the case of multi-domain proteins, only the catalytic cytoplasmic domains of the proteins were considered. The best available structure from the wwPDB database (wwPDB consortium, 2019), the ideal biological assembly, as well as the relevant chain (in multimeric structures) were selected based on resolution and missing parts. Canonical co-factors for structures were established using the UniProt database, these were retained in the structure, and all other ligands, ions, and water molecules were removed from the structure (**[SI 1](https://docs.google.com/spreadsheets/d/1wuE5v_9v-_mS6be1go6eWHxTpCl2vfk3nP-mMJ6-f0Q/edit#gid=757186111)**). The residue indexes were mapped using the SIFTS database (Dana et al., 2018). After visual inspection of each target protein, these four key problematic regions/positions were identified: (i) missing regions, i.e., low resolution regions in the crystal structure; (ii) long, missing and/or intrinsically disordered regions not influencing the catalytic site of the protein; (iii) missing atoms in the side chain; (iv) amino acids requiring identity correction, i.e., the sequence in the 3D structure did not correspond to that recorded in UniProt.

Each protein structure which required any of these structural improvements (for the aforementioned problematic regions/positions i, iii, or iv), was modelled using MODELLER version 9.24, 2020/04/06, r11614 (Eswar et al., 2008). The modelling was guided by the UniProt-PDB alignment provided by SIFTS. Regions identified as intrinsically disordered (repair ii) were omitted from the modelling. Custom extensions of three MODELLER python classes (Environment, Model, and AutoModel) were developed to ensure that the produced models: (i) incorporated any relevant co-factor from the template, (ii) were not optimised on the regions that did not require repairs, and (iii) structures containing multiple chains could be modelled and minimised at once. If no experimental structure was available, the AlphaFold database (Jumper et al., 2021) was searched. The mutant structure was generated by introducing the desired mutation in the target wild type structure by MODELLER, and it was guided by a trivial alignment between the wild type and the mutant sequences.

For each protein structure, inconsistent torsion angles, total energy, or Van der Waals clashes were reduced using RepairPDB feature of FoldX 4.0 (Blanco et al., 2018). Then minimization of structures was performed in Rosetta 3.11-static (Kellogg et al., 2010) with constraints using the Talaris2014 force field (O’Meara et al., 2015). The wild type and mutant structures were then aligned using DeepAlign 1.135-2-foss-2018b (Jiménez-Moreno et al., 2021) to ensure that their coordinates match for further analysis.

### Protein Stability Prediction

The impact of the missense mutation on the stability of the protein structure was calculated using Rosetta and FoldX. For FoldX the PssmStability command was used, water molecules were only taken from the ‘crystal’, pH was set to 7 and the number of runs was set to 5. Rosetta calculations were made on the minimised structures using the ddg_monomer command, following protocol 3 (Kellogg et al., 2010), for which the extent of sidechain repacking is set to within 8Å while using the soft-rep energy function and the Talaris2014 force field.

### Protein Function Prediction, Phylogenetic Analysis and Consensus Classification

Additionally, PropKa 3.4.0 (Rostkowski et al., 2011) was used to predict the impact of the mutation on the pKA values of the proteins, using the propka3 command. Homologous sequences with sufficient identity (more than 50%) and coverage (± 20% of the query sequence), i.e., UniRef50 sequences, were downloaded from the UniRef database (Suzek et al., 2014) and multiple sequence alignment were generated using Clustal-Omega (Sievers et al., 2011) tool from the EMBL-EBI web server (Madeira et al., 2022). This was used for conservation analysis using Jensen-Shannon Divergence algorithm (Capra & Singh, 2007) and transformed to mutability grades by using HotSpot Wizard (Sumbalova et al., 2018) thresholding. The mutations were also submitted to the HOPE (Venselaar et al., 2010) web server to collect information from a multitude of information sources; including calculations on the 3D coordinates of the protein, sequence annotations from the UniProt database, and predictions by DAS (Distributed Annotation System) services (Prlić et al., 2007). Furthermore, PredictSNP (Bendl et al., 2014), was used to predict the effect of the amino acid substitution on the target protein function through consensus classification.

### Pocket Analysis and Virtual Screening

Potential binding pockets within the structures of the analysed proteins were calculated using the prank predict command in P2Rank 2.3 (Krivák & Hoksza, 2018), the resulting pockets were visually analysed and manually optimised to cover the entire binding sites. Selected pockets were listed in **SI 2** according to their colour codes.

Virtual screening was performed on both the wild type and the mutant protein structure. A set of 4380 small molecules that were approved by the Food and Drug Administration and European Medicines Agency was taken from the ZINC database (Irwin et al., 2012). AutoDock Vina 1.1.2 (Trott & Olson, 2009) was run using the standard vina command, within a parameterized grid within each protein. The grid coordinates (**[SI 1](https://docs.google.com/spreadsheets/d/1wuE5v_9v-_mS6be1go6eWHxTpCl2vfk3nP-mMJ6-f0Q/edit#gid=757186111)**) were created by visually placing the grid on the protein structure in PyMOL using the ADPlugin (Seeliger & de Groot, 2010) and ensuring that the binding pockets with essential residues were completely within the grid. The values for the binding energy of each small molecule to a wild type structure as well as its mutant structure were used to calculate the impact of the mutation on the binding energy.

### Machine Learning Predictor Development

The predictive part of the pipeline is a machine-learning based tool that was trained on 1073 single-point mutants whose effect was classified as Oncogenic or Benign. The variants for the Benign class were selected from the gnomAD and ClinVar (Landrum et al., 2017) databases. Variants with >1% population frequency in gnomAD, variants annotated as „benign” or „likely benign” in the ClinVar database, and variants without ClinVar annotation, for which the classification as „benign” or „likely benign” is at the same time supported by applicable ACMG criteria (Richards et al., 2015) were utilised. The variants for the Oncogenic class were collected in expert-curated precision oncology knowledgebases, mainly, but not limited to,precision oncology knowledge base OncoKB by Memorial Sloan Kettering Cancer Center (Chakravarty et al., 2017), as well as The JAX Clinical Knowledgebase by The Jackson Laboratory (Patterson et al., 2016), Personalized Cancer Therapy Knowledge Base by MD Anderson Cancer Center (Kurnit et al., 2017), cBioPortal (Gao et al., 2013), and the DoCM database (Ainscough et al., 2016). Variants with conflicting interpretations across multiple sources were not included in the list. Both subsets were manually filtered for any possible overlaps with the datasets used in the PredictSNP consensus predictor and its constituents.

The entire dataset (SEQ: 509 oncogenic and 564 benign data points) was further annotated by the pipeline of PredictONCO. The following six features were calculated regardless of the structural information available: essentiality of the mutated residue (yes -1 /no - 0), the conservation of the position (the conservation grade and MSA score), the domain where the mutation is located (“cytoplasmic”, “extracellular”, “transmembrane”, “other” - one-hot encoded), the PredictSNP score, and the number of essential residues in the protein. For approximately half of the data (STR: 377 oncogenic and 76 benign data points), the structural information was available, and six more features were calculated: FoldX and Rosetta ddg_monomer scores, whether the residue is in the ligand-binding pocket obtained from P2Rank (yes -1 /no - 0), and the pKa changes of essential residues obtained from PROPKA3. The dataset is available at https://zenodo.org/records/10013764.

For the training protocol, 20% of the data in each of the two sets was kept aside for testing, chosen randomly but grouped by positions to ensure that no specific position in a protein from the test set appears in the training set. The following types of predictors were tested: the support vector machine (SVM), decision tree (DT), and XGBoost classifier (XGB), taken as they are implemented in the scikit-learn 1.2.0 and xgboost 1.7.3 libraries for Python 3.8.15. We also used the PredictSNP score alone as a baseline. For each method, we tested a set of hyperparameters based on 5-fold cross-validation implemented on the training data and receiver operating characteristic (ROC) area under the curve (AUC) as the metric (Table S1 in **[SI 3](https://docs.google.com/document/d/1QJoPHS7ms7qd6uswId3wR6TWb7iLBtvv/edit?rtpof=true&sd=true)**).

The final evaluation consisted of constructing the ROC and Precision-Recall curves. Furthermore, a round of 100 random-state re-initialisations with different random seeds was performed to evaluate the robustness of the final models. For the area under the ROC curve and the average precision values, we reported the average and standard deviation from the 100 runs. Since any change to the predictor or data split results in a different set of x-axis coordinates in the ROC and Precision-Recall curves, we used a fixed grid of 30 points and applied 1D linear interpolation to obtain the y-axis value for each iteration. These values were then plotted as 10% and 90% quantiles.

The version numbers for softwares and python packages that were used are noted in SI 4.

## Results

### Development of a Fully Automated Computational Workflow

We created a bioinformatics pipeline for structure and sequence based analysis of the effects of missense mutations on cancer-related proteins (Figure S1). Since the pipeline requires curated protein structures, a method for curation was developed and applied to a list of 44 proteins (**SI 1**) which were then tested to ensure they can be handled in the pipeline. The pipeline was assembled using multiple bioinformatics tools, databases, and techniques. **Figure 1** represents a schematic outline of the pipeline, the output of which ultimately feeds into the machine learning predictor. The predictor gives a binary decision on the effect of mutation with confidence score which is helpful in the summation and comprehension of results. Three cases of oncological interest were then studied using the developed method.

**Figure 1.**
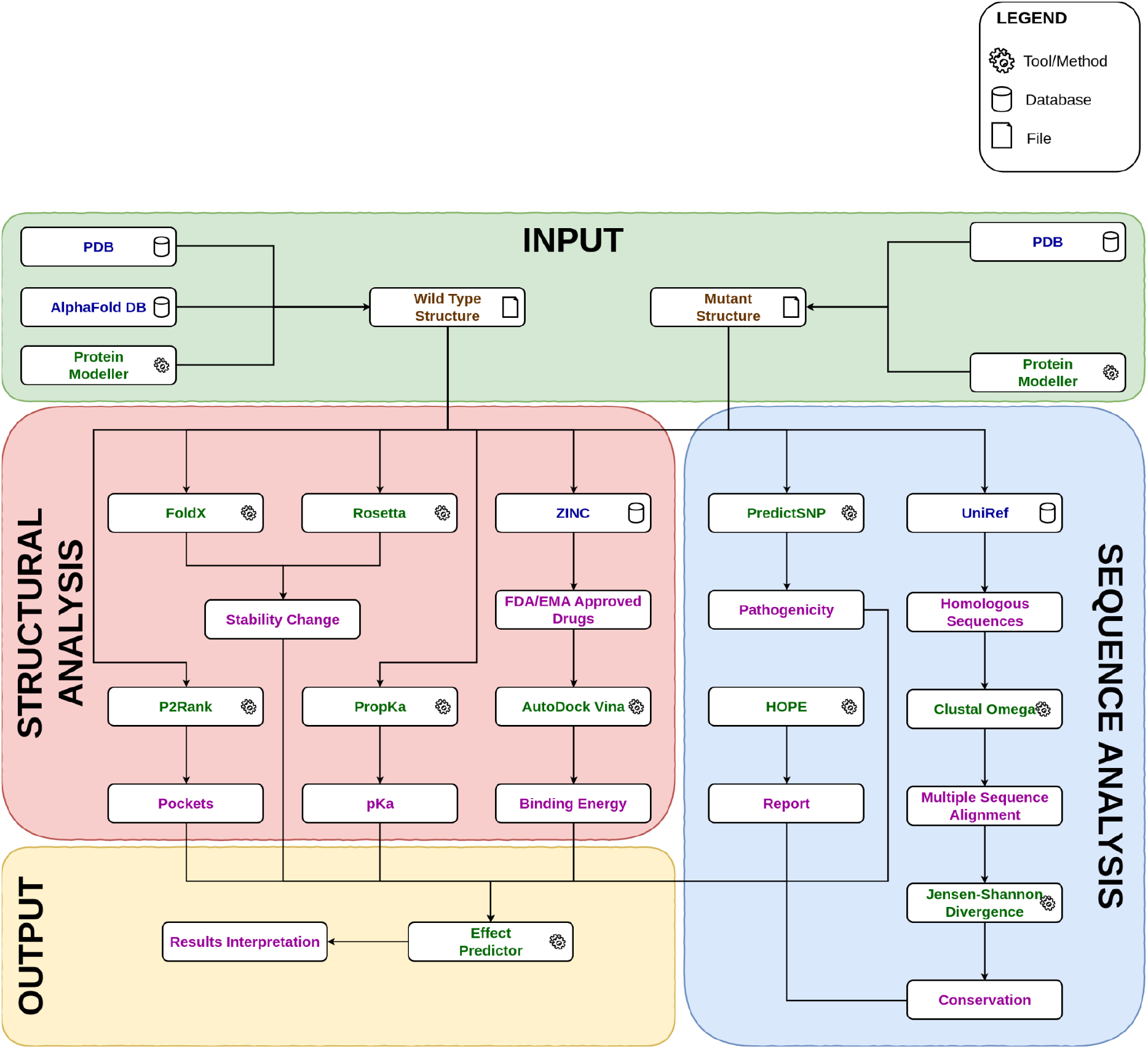
A schematic representation of the bioinformatic pipeline used to predict the effect of a missense mutation on the oncogenicity of the protein.

### Training of Sequence-Based and Structure-Based Machine Learning Predictors

Initially, we trained three different types of predictors, covering different trade-offs between explainability and flexibility, and compared their performance with the baseline model using the PredictSNP score alone. After optimising the hyperparameters (**Table S1** in **[SI 3](https://docs.google.com/document/d/1QJoPHS7ms7qd6uswId3wR6TWb7iLBtvv/edit?rtpof=true&sd=true)**), we evaluated the performance on the held-out 20% of the dataset split by position in a protein. The support vector machines and XGBoost classifiers showed superior yet similar performance based on the area under the ROC curve and the average precision from the Precision-Recall curve (**Figure 2**), also confirmed statistically (**Figure S2** in **[SI 3](https://docs.google.com/document/d/1QJoPHS7ms7qd6uswId3wR6TWb7iLBtvv/edit?rtpof=true&sd=true)**). We selected the XGBoost predictor for the final model due to the interpretability of its scores: the SVM model evaluation is based on the signed distance to the separating hyperplane, without intuitive interpretation. On the other hand, the XGBoost classifier directly returns the probability that a particular mutation is oncogenic. The final XGBoost predictor is made up of 15 and 9 decision trees of the depth of 1 for structure and sequence data sets, respectively. The feature importance scores revealed that the PredictSNP score and conservation had the highest information gains (**Figure S3** in **[SI 3](https://docs.google.com/document/d/1QJoPHS7ms7qd6uswId3wR6TWb7iLBtvv/edit?rtpof=true&sd=true)**). We also tested if using the train/test split by proteins would compromise the performance and saw only a marginal decrease (Figure S4 in **[SI 3](https://docs.google.com/document/d/1QJoPHS7ms7qd6uswId3wR6TWb7iLBtvv/edit?rtpof=true&sd=true)**), indicating the significant potential of the pipeline for other protein targets. The balanced accuracy for the sequence-based XGBoost predictor is 87%, and for the structure-based XGBoost predictor is 90%.

**Figure 2.**
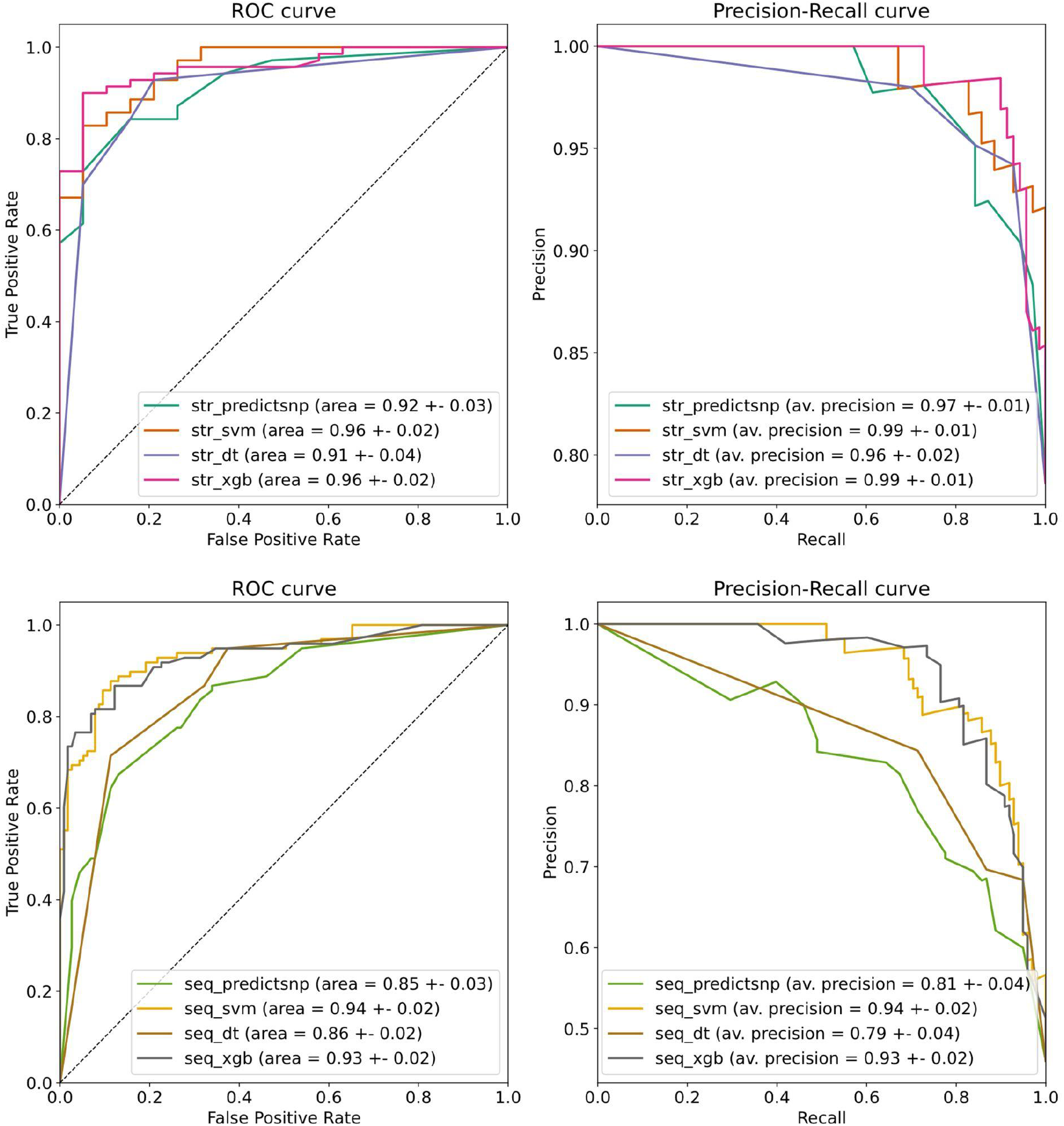
The Receiver Operating Characteristic and Precision-Recall curves based on held-out test sets. Top: classifiers trained on the dataset with the structural features available (STR). Bottom: classifiers trained on the dataset with the sequence-only features (SEQ). Both the support vector machine (SVM) and XGBoost (XGB) showed comparable performance superior to the baseline model and decision tree (DT). The reported errors are standard deviations obtained by bootstrapping (N=1000). The PredictSNP score was used as the baseline.

We also compared the performance of our predictor on the test set againsts several other models (Table 1). We evaluated the following individual scores as baselines: conservation, predictSNP, FoldX, and Rosetta. In addition, we evaluated the performance of the ESM variants model, a recently published workflow based on the 650-million-parameter protein language ESM1b, which was used to score all possible missense variant effects in the human genome (Brandes et al., 2023). In both settings (SEQ and STR), PredictONCO showed superior performance.

**Table 1.** Comparison of PredictONCO with other models on the test set. The models selected for comparison were individual features and the ESM variants predictor. The reported errors are standard deviations obtained by bootstrapping (N=1000).

|  | Predictor | ROC AUC↑ | Avg. Precision↑ |
| --- | --- | --- | --- |
| SEQ | **PredictONCO** | **0.932 ± 0.018** | **0.934 ± 0.018** |
|  | conservation | 0.872 ± 0.026 | 0.802 ± 0.042 |
|  | predictSNP | 0.845 ± 0.027 | 0.808 ± 0.041 |
|  | ESM variants | 0.923 ± 0.018 | 0.911 ± 0.023 |
| STR | **PredictONCO** | **0.955 ± 0.020** | **0.988 ± 0.006** |
|  | FoldX | 0.575 ± 0.064 | 0.867 ± 0.037 |
|  | Rosetta | 0.628 ± 0.064 | 0.876 ± 0.039 |
|  | conservation | 0.937 ± 0.037 | 0.970 ± 0.020 |
|  | predictSNP | 0.918 ± 0.030 | 0.973 ± 0.011 |
|  | ESM variants | 0.929 ± 0.027 | 0.981 ± 0.009 |

### Case Studies with Selected Cancer-associated Proteins

The following case studies demonstrate scenarios in which the tool has helped to facilitate further clinical decision-making. The respective variants featured in the case studies were identified across research projects utilizing high-throughput DNA sequencing techniques, which were conducted by the co-authors of this manuscript.

### Case study 1 - Platelet Derived Growth Factor Receptor Beta PDGFRB N666T

In a patient with myofibroma, sequencing analysis revealed an N666T variant of the PDGFRB protein (UniProt ID:P09619). Even though some mutations of the N666 residue, including N666K (Iwamura et al. 2023), N666H (Pond et al. 2018), or N666S (Ortiz et al. 2020), have already been documented in myofibroma patients, N666T, in particular, lacks published functional evidence and was reported in a total of one patient in combination with another mutation. Therefore, a comprehensive assessment of its effect would provide further confirmatory evidence on the variant’s pathogenicity, which is substantial, given the therapeutic implications of receptor tyrosine kinase inhibition. Conservation status showed high evolutionary conservation, which is typically damaging to the protein. For amino acid 826, one of the essential catalytic residues, a large increase in dissociation constant was predicted, suggesting a significant functional impact. Both stability predictors suggested a deleterious effect, which is also in agreement with the deleterious effect on protein function predicted by PredictSNP. Given all this data, the oncogenic effect was predicted by the XGBoost classifier with 100% confidence. Furthermore, in virtual screening, Sunitinib showed a slightly better increase in binding affinity compared to Imatinib, which was used as a drug of choice in different myofibroma preclinical studies, making Sunitinib a suitable alternative option for therapeutic planning. The full report can be accessed at - https://loschmidt.chemi.muni.cz/predictonco/job/pdgfrb_N666T

### Case study 2 - Angiopoietin-1 receptor TIE2 G1036D

In a patient with a vascular tumour, sequencing analysis revealed a G1036D variant in the TIE2 (UniProt ID:Q02763) gene. The G1036D variant represents a previously undescribed alteration, which has not been documented in the literature, clinical, or population databases of genetic variants. Given the rapidly evolving field of vascular tumour genetics and the possibility of targeted therapeutics administration, identifying novel potentially activating alterations is vastly important. Although the residue is non-essential, moderately evolutionarily conserved, and only moderate changes were predicted for the catalytic residues, the overall impact was evaluated by the XGBoost classifier as oncogenic with a 99% confidence score and was based on a deleterious prediction by both the PredictSNP algorithm and stability predictors FoldX and Rosetta. This could be approached as a basis to facilitate further functional tests to measure mutant receptor phosphorylation and, if proven as activating, introduce a considerable therapeutic opportunity (by potentially using one of the suggested inhibitive compounds such as Ecteinascidin, Ponatinib, etc., or other inhibitors of downstream signalling cascade) as well as an addition to the knowledge on disease pathogenesis. The full report can be accessed at - https://loschmidt.chemi.muni.cz/predictonco/job/tie2_G1036D

### Case study 3 - Cellular Tumor Antigen p53 K101Q

In next-generation sequencing screening for cancer predispositions, the K101Q variant of p53 (UniProt ID:P04637) was identified in an individual with a negative family history of cancer. p53 represents the most commonly altered gene in all cancers, and p53 variants predispose to cancer development when of germline origin. Therefore, a careful assessment must be performed for further genetic counselling. The respective variant has not been documented in the literature or functionally characterised. With lacking evidence from literature and databases of genetic variants, typically only prediction algorithms that employ sequence-based information without structural data are available. Therefore, combining both structural and sequence related perspectives might yield a more accurate prediction. The XGBoost classifier predicted the mutation as neutral with an 81% confidence score, supported by both the PredictSNP prediction and the stability predictors. Information on evolutionary conservation showed that the wild-type residue is not conserved at this position, which may suggest that the variant is not damaging to the protein. Based on these results and no family history of cancer, the variant should not influence subsequent clinical management. Given the importance of p53 variants in both somatic and germline contexts and their same functional impact, this case study exemplifies the utility of the tool in the assessment of hereditary cancer predisposition. The full report can be accessed at the following link - https://loschmidt.chemi.muni.cz/predictonco/job/p53_K101Q

## Discussion

Prediction of the effect of missense mutations on cancer-related protein structures is a complicated task. This paper presents our pipeline for tackling this problem thus allowing clinical bioinformaticians to easily run multiple cancer-related analyses for their target mutations on a curated list of proteins.

A major part of the pipeline capitalises on structural bioinformatics and it requires the presence of good quality protein structures for accurate analysis. However, a good amount of cancer-associated structures are transmembrane channels and thus only have fragmented domain level structures. Some of them can be multimeric, and thus modelling proves a challenge. Despite AlphaFold (Jumper et al., 2021) being touted as a major groundbreaker in the field of protein structure modelling, it proves inefficient in modelling large multi-subunit, multimeric proteins as quaternary domain level interactions are difficult to model. Thus the structural bioinformatics part of the pipeline is limited to working with high-quality structures at the domain level. AlphaFold-Multimer (Evans et al., 2021) can be used to predict the multimeric conformation in 70% of heteromeric cases and 72% of homomeric cases to limit this problem; it is unclear whether this accuracy of predictions is viable for working with oncogenic or tumour suppressor proteins, especially when the final prediction will likely be used in a medical context.

Currently, the web server provides predictions for 44 target proteins, which were selected based on their relevance to the field of oncology. Appropriate processing of a new structure to be used in the pipeline requires expert level knowledge of multiple bioinformatic tools. Curation in this field is a recognized bottleneck, especially in the case of the interpretation of results (Bungartz et al., 2018.) The addition of new target proteins to the internal database connected to the PredictONCO web server is possible and it is offered to the user community based on direct requests. Once a protein is curated, all mutations on its structure can be easily analysed. Moreover, the pipeline can also work with sequence only data and the trained XGBoost classifier can also reliably predict using only the sequence based features with only a 4% drop in average precision.

The pipeline has no standard run time as it mostly depends on whether structural analysis needs to be performed along with sequence-based analysis or not. The structural analysis increases the computational load and the complexity of the structure can further increase the run time. However, the calculations generally do not take more than two days to complete. It is unclear whether this time frame is long or short as run time benchmarking would require the existence of other similar tools, techniques or pipelines for comparative purposes, and specialised methodologies that deal with the same case do not exist. However, this time window meets the initial requirements for the use of the web server in clinical practice as well as for research and educational purposes. Furthermore, it helps assist in making the result interpretation step easier as interpretation itself is a recognized bottleneck (Bungartz et al. 2018.)

Comparison to other similar tools is difficult as, as of this writing, we did not come across a pipeline integrating multiple approaches to predict the effect of a missense mutation on a cancer-related protein. However several databases and online data integrating tools do exist. The two most prominent of these databases would be the International Cancer Genome Consortium (ICGC) (The International Cancer Genome Consortium, 2010) and The Cancer Genome Atlas (TCGA) (Weinstein et al. 2013). Furthermore, survival analysis tools also exist and are primarily based on 4 types of data: (i) mRNA data, such as PRECOG (Gentles et al., 2015), (ii) ncRNA data, such as OncoLnc (Anaya, 2016), (iii) DNA methylation and mutation data, such as cBioPortal (Gao et al., 2013), and (iv) Protein data, such as TCPA (Li et al., 2013). Additionally, the Swiss-PO web tool for mapping gene mutations on the 3D structure can be used, but it only allows for intuitive and qualitative analysis of mutations that have already been experimentally determined (Krebs et al., 2021). Comparison to the aforementioned database PSnpBind is also difficult as it only catalogues changes to binding affinities of ligands due to binding site single-nucleotide polymorphisms (SNPs) (Ammar et al., 2022).

Our pipeline currently only supports missense mutations, as it is unable to handle insertions, deletions, or fusions of oncogenic proteins because individual tools in the pipeline are not able to analyse them. However, substitutions do make up a large number of cancer associated mutations as a large number of genes associated with various cancer types contain single nucleotide variants (Deng et al., 2017). For common solid tumours, 95% of cancer driver mutations in humans are single-base substitutions. Approximately, 90.7% of these result in the amino acid being substituted for another, non-synonymous one (Darbyshire et al. 2019). Thus, even though insertions, deletions, and fusions cannot be analysed using the pipeline, it still provides predictions for a significant majority of cancer-related alterations. The tool is freely accessible to the community of bioinformaticians and medical doctors and will provide fast and useful analysis of data from sequencing of patient samples.

## Availability of data and materials

- The pipeline is available as a web server, at https://loschmidt.chemi.muni.cz/predictonco/.
- The list of proteins, definition of binding pockets, and ML model validation are attached as supplementary files.
- The training and testing dataset are available at https://zenodo.org/records/10013764.

