## Supplementary material for "A Computational Workflow for Analysis of Missense Mutations in Precision Oncology": SI 3

**Supplementary Information 3**

**Table 1.** The set of hyperparameters tested and the final values selected for further analysis. The optimal hyperparameters were selected based on 5-fold cross-validation and ROC AUC metric.

| **Method** | **Hyperparameters optimized** | **Values selected (STR)** | **Values selected (SEQ)** |
| --- | --- | --- | --- |
| SVM | penalty: {'l1','l2'}  kernel: {‘linear’, ‘poly’, ‘rbf’, ‘sigmoid’} | penalty: 'l1'  kernel: ‘linear’ | penalty: 'l1'  kernel: ‘linear’ |
| DT | max_depth: 1..10  min_samples_split: 2..10 | max_depth: 2  min_samples_split: 2 | max_depth: 2  min_samples_split: 2 |
| XGB | n_estimators: 1..40  max_depth: 1, 2, 3  learning_rate: 0.001..1000 | n_estimators: 15  max_depth: 1  learning_rate: 1 | n_estimators: 9  max_depth: 1  learning_rate: 1 |

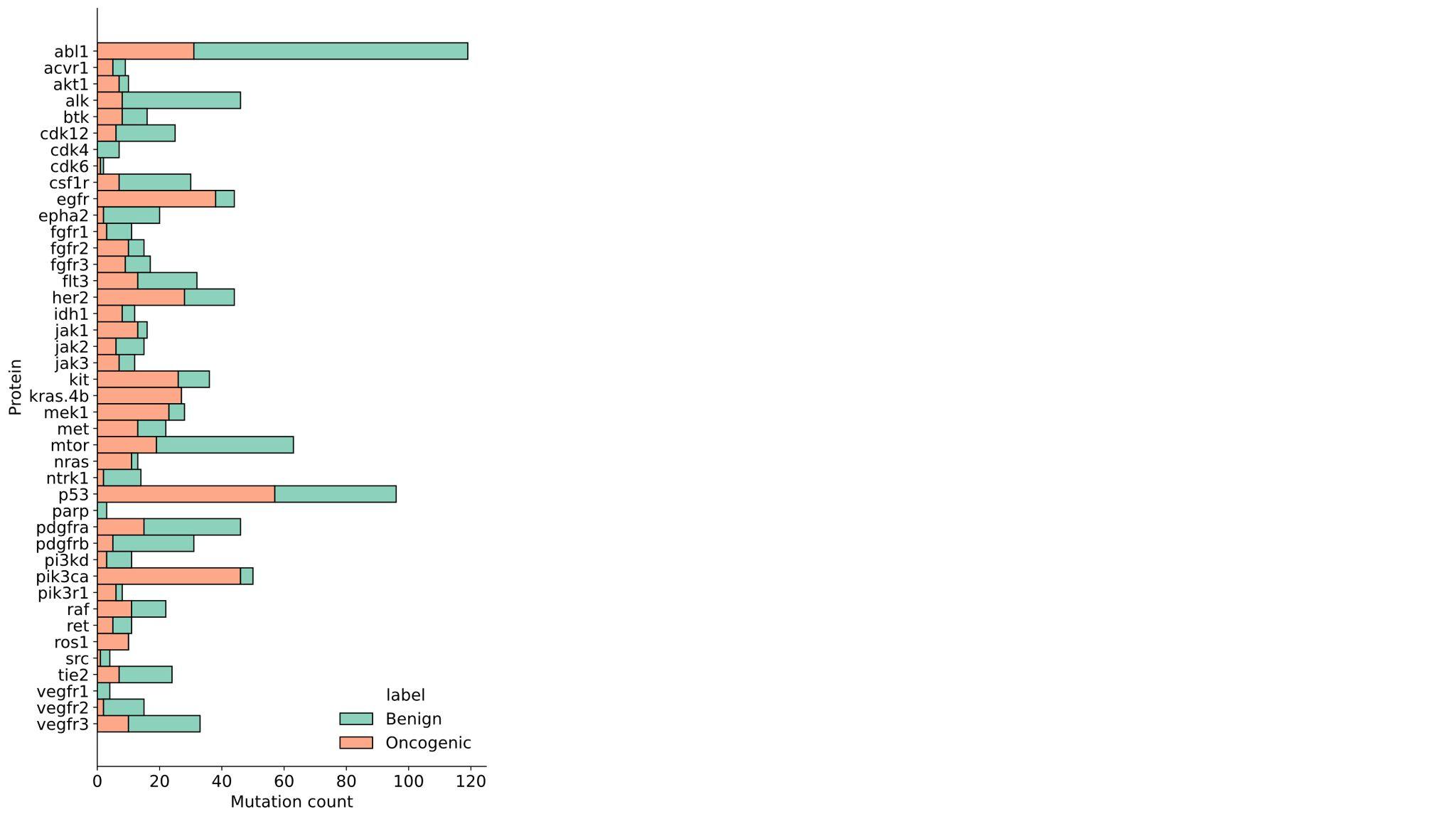

**Figure S1.** **The distribution of mutations by protein in the entire training dataset.** Most proteins contributed around 10-50 variants to the dataset.

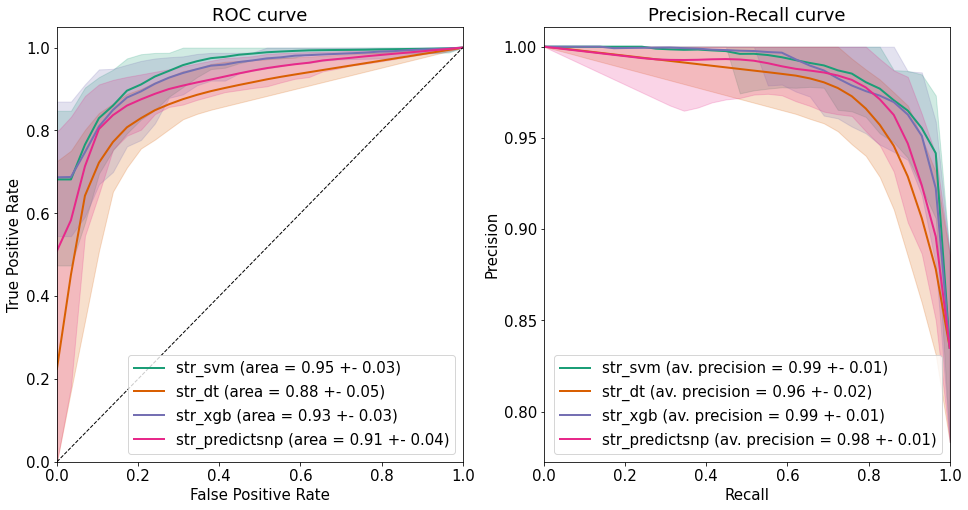

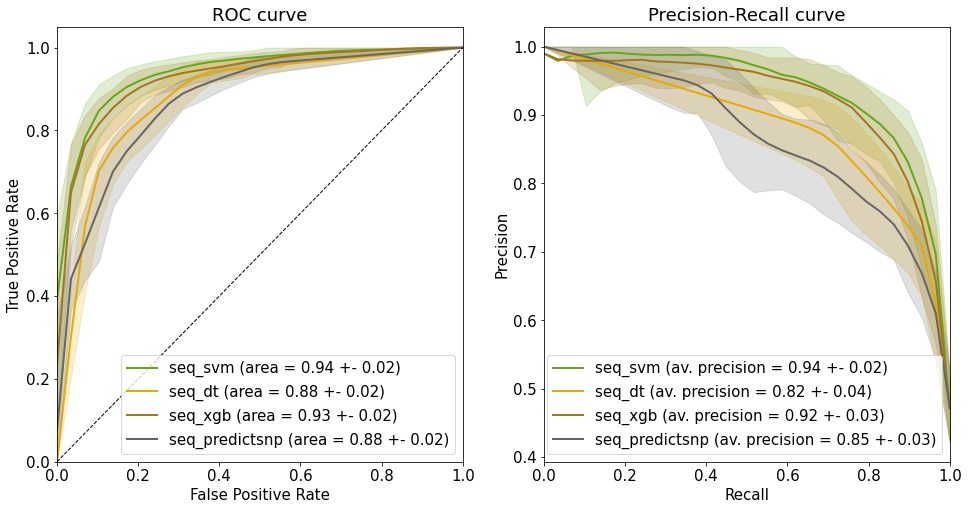

**Figure S2.** **Resampling of ROC and Precision-Recall curves 100 times**. The shaded area denotes the 10% and 90% quantiles for the y-axis values, linearly interpolated at fixed grid points of each curve. The area under the ROC curve and average precision are reported as means and standard deviations. **Top**: classifiers trained on the dataset with the structural features available (STR). **Bottom**: classifiers trained on the dataset with the sequence-only features (SEQ).

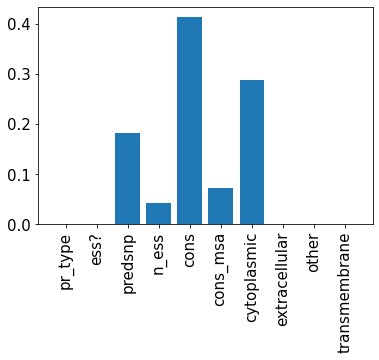

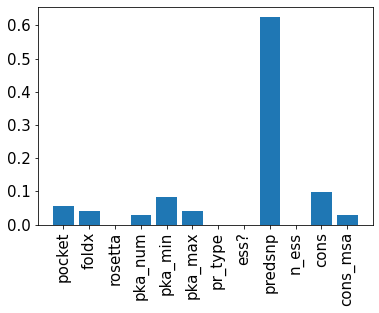

**Figure S3.** **Feature importance scores for the final STR (left) and SEQ (right) XGBoost classifiers.** The two features with the largest contribution to the information gain are the predictSNP score and conservation.

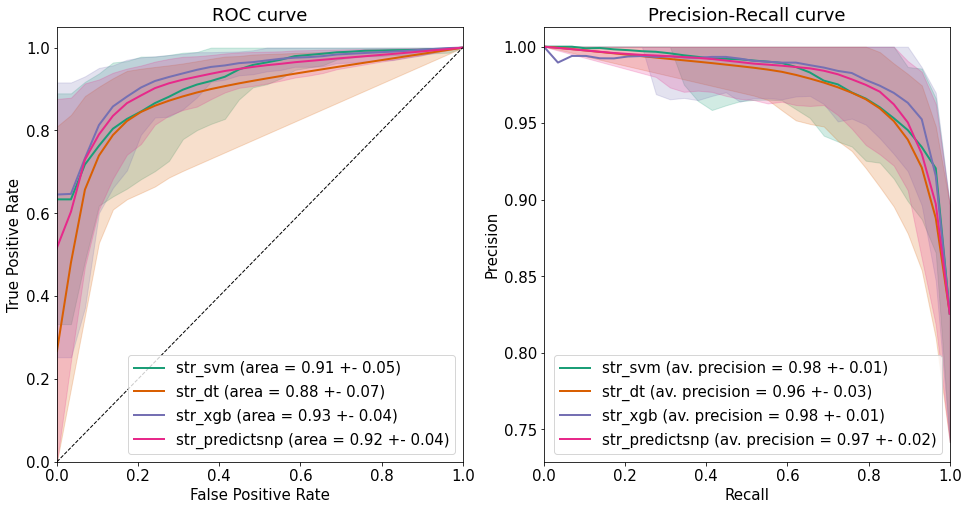

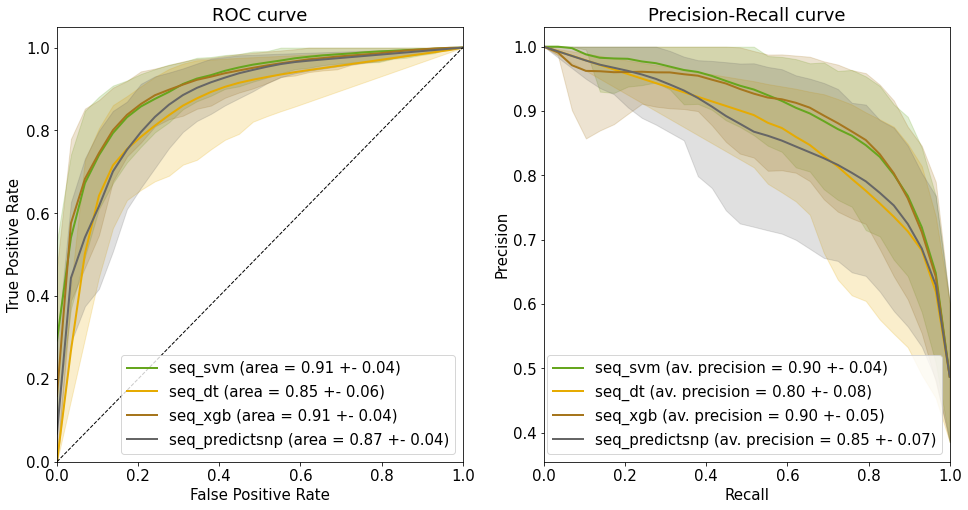

**Figure S4.** **Resampling of ROC and Precision-Recall curves for the predictors with the split based on the protein.** The shaded area denotes the 10% and 90% quantiles for the y-axis values, linearly interpolated at fixed grid points of each curve. The area under the ROC curve and average precision are reported as means and standard deviations.
