## Supplementary material for "A Computational Workflow for Analysis of Missense Mutations in Precision Oncology": SI 4

**Supplementary Information 4**

MODELLER 9.24, 2020/04/06, r11614

FoldX: 4.0

DeepAlign: 1.135-2-foss-2018b

Rosetta: 3.11-static

p2rank: 2.3

propka: 3.4.0

AutoDock Vina: 1.1.2

numpy=1.23.5

scipy=1.9.3

xgboost=1.7.3

scikit-learn=1.2.0
